# Polyphasic discrimination of *Shewanella seohaensis* from closely related species and a whole-genome multilocus (wgMLST) scheme for the evaluation of diversity within this *Shewanella* clade

**DOI:** 10.1101/2025.02.24.639908

**Authors:** Maria de Oliveira Firmino, Mykyta Forofontov, Ricardo O. Louro, Alberto J. Martín-Rodríguez, Mário Ramirez, Catarina M. Paquete

**Affiliations:** Instituto de Tecnologia Química e Biológica António Xavier, Universidade Nova de Lisboa, Av. da República, 2780 – 157 Oeiras, Portugal; Instituto de Microbiologia da Faculdade de Medicina da Universidade de Lisboa, Avenida Prof. Egas Moniz, P-1649-028 Lisboa, Portugal; Department of Clinical Sciences, University of Las Palmas de Gran Canaria, Paseo Blas Cabrera Felipe, s/n, 35016, Las Palmas de Gran Canaria, Spain; Department of Microbiology, Tumor and Cell Biology, Karolinska Institutet, Solnavägen 9, 171 65 Solna, Sweden

## Abstract

*Shewanella* is an environmentally ubiquitous genus with significant roles in bioelectrochemical applications and human infections. However, identification problems involving *Shewanella putrefaciens*, *Shewanella xiamenensis,* and *Shewanella seohaensis*, have been reported, potentially hindering research progress in these areas. In this study, we explored how to discriminate between these species. By comparing the genomes of *Shewanella* spp. available in public databases with that of the newly sequenced strain DSM9451, we showed that this strain is a member of the species *S. seohaensis*. Of the eight public genomes associated with this species, only two were correctly identified in public databases. Phenotypic analysis revealed distinct features of *S. seohaensis* with respect to *S. putrefaciens* and *Shewanella decolorationis.* However, only differences in the intensity of biochemical reactions were observed between *S. xiamenensis* and *S. seohaensis.* To discriminate between these closely related species and explore their diversity, a whole-genome multilocus sequence typing scheme was developed. The scheme distinguished related species and revealed significant diversity among *S. seohaensis* and *S. xiamenensis* isolates, suggesting that both species may harbor isolates with significantly different metabolic properties. These differences are so wide within the clade that may explain why these two species are difficult to distinguish. The identification of exclusive genes of each species allowed the design of a simple molecular method to differentiate *S. seohaensis* from closely related species, which will help in clarifying its role in human infections and environmental processes.

**Importance:** Misidentification within the *Shewanella* genus, particularly between *S. putrefaciens*, *S. seohaensis,* and *S. xiamenensis*, has been reported and misperceives scientific research on *Shewanella spp*. in diverse fields, including both biotechnological applications and human infections. Near-complete 16sRNA gene sequencing fails to correctly classify many *Shewanella* species, and MALDI-TOF systems used in clinical microbiology laboratories are suboptimal species-level identification. In this study, we identified phenotypic characteristics that can guide differentiation and classification, and building upon the identification of species-specific genes, we suggest an accurate and cost-effective molecular test as an alternative to genome sequencing. The proposed wgMLST scheme allows the exploration of species and strain diversity, highlighting the limitations of generalizing results from studies of a single strain. As an emergent pathogen and biotechnological candidate, the proper identification by a single molecular test will enhance the insights about these species towards biotechnology development and public health safety.

## Introduction

The genus *Shewanella,* proposed by MacDonell and Cowell in 1985 (1), is a ubiquitous and diversified genus that belongs to the class of Gammaproteobacteria, order Alteromonadales, and family *Shewanellaceae* (2). The first isolate reported from this genus had several classifications, including *Achromobacter putrefaciens*, *Pseudomonas putrefaciens,* and *Alteromonas putrefaciens,* until its final classification as *Shewanella putrefaciens* (3). *Shewanella* spp. can be found in fresh and seawater and human samples (4). In recent years, the interest in this genus has increased due to its importance in bioremediation, microbial electrochemical technologies, and biosensing (5) and by being identified as a potential emerging pathogen (6–8). All human isolates reported until 1995 were mistakenly identified as either *Pseudomonas putrefaciens* or *Alteromonas putrefaciens* (9). *Shewanella algae* was recognized as a pathogen in the early 1990s, being easily distinguished due to its distinctive phenotypic differences, such as the ability to grow at higher temperatures and salt concentrations (3, 10). It was only after the first DNA sequencing of several isolates of the *Shewanella* genus in 2002 (11) that other *Shewanella* species, including *Shewanella xiamenensis*, were identified as pathogenic (12). The identification of other *Shewanella* species in samples from human patients, such as *Shewanella carassii* (13), suggests that the breadth of potentially pathogenic species within this genus might be wider than initially thought. Improved identification systems and changing environmental conditions (such as climate change and socio-geographic distribution) (8, 14) are contributing to a rise in documented cases of *Shewanella* infections (15). *Shewanella* spp. are frequently present in polymicrobial infections (3, 8) and are considered important environmental reservoirs of antimicrobial resistance genes (16).

Genome sequencing stands out as the most suitable method to achieve accurate identification within the *Shewanella* genus. To resolve the taxonomy of the genus *Shewanella,* Martín-Rodríguez and Meier-Kolthoff sequenced all *Shewanella* type strains that lacked information in public repositories and reconstructed a genus-wide phylogeny with all sequenced *Shewanella* strains at that time (17), demonstrating that relying solely on 16S RNA sequence-based identification has limitations within this genus. Given that sequencing of new isolates is a laborious and expensive identification method, the issue of misidentification of *Shewanella* species persists. This problem is particularly evident in infection-associated *Shewanella* isolates (16, 18, 19), as standard clinical microbiology identification methods, frequently carried out through automated or semi-automated methods, often rely on incomplete or inaccurate databases (20). A notable example is the systematic difficulty in correctly identifying *Shewanella seohaensis, S. xiamenensis,* and *S. putrefaciens* (21). Thorell and colleagues demonstrated the existence of misidentified isolates, including *S. putrefaciens* SA70 and *S. putrefaciens* NCTC12093 (18), while the work of Martín-Rodríguez and Meier-Kolthoff determined that these isolates belong to the species *S. seohaensis* (17).

In this study, we focus on differentiating *S. seohaensis*, *S. xiamenensis,* and *S. putrefaciens* by phenotypic characteristics and molecular methods. Towards this, the phylogenetic classification and phenotypic characteristics of two pathogenic *Shewanella* strains, *S. putrefaciens* DSM9451 and *S. xiamenensis* (HI32665) isolated from human clinical samples, were determined. This research enhances our current understanding of *Shewanella* taxonomy and also investigates the differences between *S. seohaensis* and closely related species. It provides the tools for their differentiation in laboratories lacking genome sequencing capabilities, which potentially helps to clarify the role of this species in human infections and environmental processes. Additionally, this study proposes a whole-genome multilocus sequence typing (wgMLST) scheme to help differentiate these species and better understand their population structure.

## Results and Discussion

### *Shewanella putrefaciens* DSM9451 is a member of the *Shewanella seohaensis* species

The type strain of *S. xiamenensis* was first isolated from a sediment sample of the coastal area of Xiamen, China (22). This species was also isolated from human infections (13, 23), establishing it as one of the pathogenic members of the *Shewanella* genus. The *S. xiamenensis* strain (HI32665) used in this work was isolated from a coproculture at a Portuguese Hospital and was initially identified as *Shewanella xiamenensis* by MALDI-TOF (MALDI Biotyper MSP Identification). The species assignment for this strain was conclusively achieved by analyzing its genome sequence relatedness with respect to the reference genome of the type strain of *S. xiamenensis*. The values of *d*DDH (74.9 % [confidence interval 71.9 – 77.7]) and ANIb (97.0%) were concordant with the identification of HI32665 as *S. xiamenensis*.

*Shewanella putrefaciens* DSM9451, initially designated as CL 256/73 and first described by Holmes *et al.* (24), was isolated from a mixed culture from the cerebrospinal fluid of a 1-year-old infant, with a suspicion of meningitis. This strain was acquired for this study from the DSMZ repository, where it is identified as *S. putrefaciens*. However, MALDI-TOF (MALDI Biotyper MSP Identification) classified this strain as *S. xiamenensis*. Distinction between these species often eludes such systems (21), since their accuracy depends on the databases supporting the analysis. To evaluate the capacity of MALDI-TOF MS to discriminate between the species *S. seohaensis*, *S. xiamenensis,* and *S. putrefaciens*, the strains *S. seohaensis* SA70, *S. putrefaciens* 95^T^ and *S. seohaensis* CCUG60900^T^ were analyzed by the MALDI Biotyper MSP Identification system. The results revealed that while the system could differentiate *S. putrefaciens,* it was unable to distinguish between *S. seohaensis* and *S. xiamenensis*, identifying strains of both species as *S. xiamenensis*. Analysis of genomic data using Kraken suggested that *S. putrefaciens* DSM9451 was indeed *Shewanella bicestrii*. However, this taxon is a non-validated species, also referred to as *Shewanella* sp. JAB-1 (15, 16). According to Martín-Rodríguez and Meier-Kolthoff (17), the affiliation of *Shewanella* sp. JAB-1 with the species *S. seohaensis* suggested a different classification for *S. putrefaciens* DSM9451, which we will further refer simply as DSM9451.

For a deeper understanding of the phylogenetic classification of DSM9451, a comprehensive phylogenetic analysis was undertaken. This analysis involved comparing the genome of DSM9451 with all 822 genomes of *Shewanella* available on the National Center for Biotechnology Information (NCBI) on the 17^th^ of June 2024 by GBDP. The results revealed that DSM9451 and *S. seohaensis* NCTC12093 have nearly identical genomes (*d*DDH = 99.9 % [99.9 - 100]) (see Figure 1). Furthermore, the comparison between DSM9451 and *S. seohaensis* CCUG6090 (*d*DDH = 70.7 % [67.7 - 73.6]), *S. seohaensis* SA70 (*d*DDH = 76.2 % [73.2 - 79]), *S. seohaensis* JAB-1 (*d*DDH = 71.9% [68.9 - 74.8]), and *S. seohaensis* BC20 (*d*DDH = 69.4% [66.4 - 72.3]) confirmed that they all belong to the same species. DSM9451 also had a *d*DDH value >70% with *Shewanella* sp. GD03713 (74.8 % [71.8 – 77.6]), indicating that *Shewanella* sp. GD03713 is of the same species as DSM9451. When compared with *S. putrefaciens* 95, the type strain of *S. putrefaciens*, the value of *d*DDH is 22.70 % [20.4 - 25.1] (Table 1). Comparing these strains with the type strain of *S. seohaensis, S. seohaensis* CCUG60900, we obtained similar results, except for *Shewanella* sp. GD03713, where the dDDH values are on the limit to consider this strain as not belonging to this species. However, the ANI results support the identification of GD03713 as *S. seohaensis* (Table 1). The ANI values presented in Table 1, reinforce that DSM9451 does not fall within the *S. putrefaciens* species.

**Figure 1.**
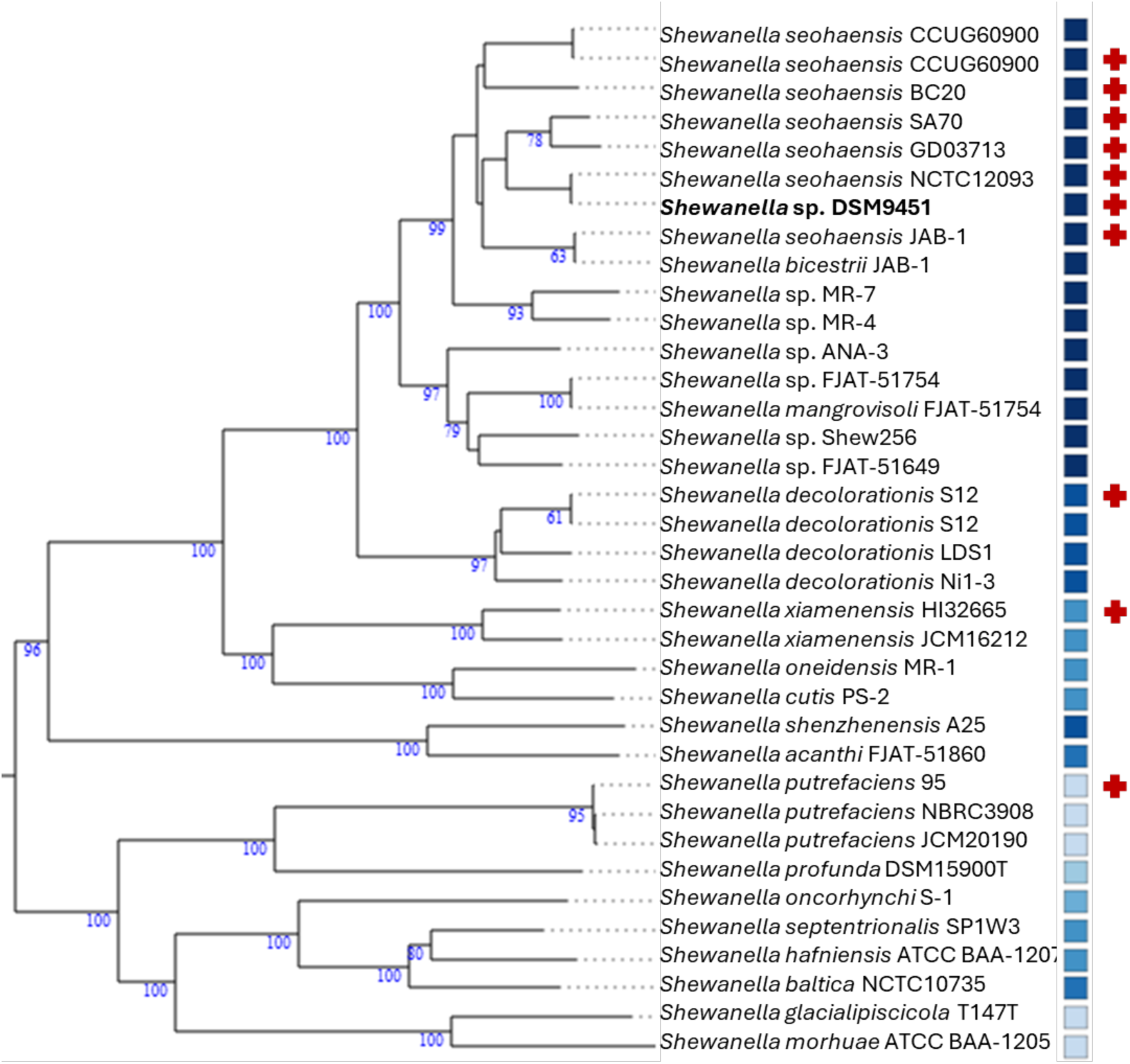
Whole-Genome-based GBDP tree. Blue shades represent the G+C percentage with darker corresponding to a higher percentage. User-provided genomes are marked by a red cross. Branch lengths are scaled in terms of GBDP distance formula d6. The numbers above branches are GBDP pseudo-bootstrap support values from 100 replicates.

**Table 1.** Matrix of dDDH and ANIb results. dDDH values cut-off is at 70% and ANI values cut-off is at 95 %. The grey shadow boxes are those that belong to the same species (the confidence interval for the estimate includes 70%). A - *Shewanella* sp. DSM9451; B - *S. seohaensis* NCTC12093; C - *S. seohaensis* JAB-1; D - *S. seohaensis* SA70; E - *S. seohaensis* CCUG60900; F - *S. seohaensis* BC20; G - *Shewanella* sp. GD03713; H - *S. putrefaciens* 95.

|  | Phylogeny analysis | A | B | C | D | E | F | G | H |
| --- | --- | --- | --- | --- | --- | --- | --- | --- | --- |
| A | dDDH % |  | **99.9**<br>[99.9 – 100] | **71.9**<br>[68.9 – 74.8] | **76.2**<br>[73.2 – 79.0] | **70.7**<br>[67.7 – 73.6] | **69.4**<br>[66.4 – 72.3] | **74.8**<br>[71.8 – 77.6] | 22.7<br>[20.4 – 25.1] |
|  | ANIb % |  | **100.00** | **96.40** | **97.07** | **96.36** | **96.11** | **96.84** | 78.69 |
| E | dDDH % | **70.7**<br>[67.7 – 73.6] | **70.7**<br>[67.7 – 73.6] | **70.6**<br>[67.6 – 73.4] | **67.4**<br>[64.4 – 70.2] |  | **71.2**<br>[68.2 – 74.0] | **66.0**<br>[63.0 – 68.8] | 22.4<br>[20.1 – 24.8] |
|  | ANIb % | **96.37** | **96.38** | **96.39** | **95.87** |  | **96.51** | **95.67** | 78.55 |

**Table 2.** Bacterial strains used in this study.

| Strain | Description | Source | Strain Reference | Genome sequencing accession number | Type strain |
| --- | --- | --- | --- | --- | --- |
| *Shewanella putrefaciens* DSM9451 | Isolated in a mixed culture from cerebrospinal fluid | German Collection of Microorganisms and Cell Cultures (DSMZ) | Holmes *et al.*, 1975 (24) | This study <sup>a</sup> | NO |
| *Shewanella seohaensis* SA70 | Isolated from the sink handle of a Pakistani hospital room | Donated by Dr. Gautam Dantas | Potter *et al.*, 2018 (19)<br>Reclassification by Martín-Rodríguez *et al* (17) | GCF_002157365.2 | NO |
| *Shewanella seohaensis* CCUG60900 | Isolated from sediments at Saemankum on the western coast of Korea | Martín-Rodríguez lab collection | Yoon *et al.*, 2012 (26) | GCF_023283775.1 | YES |
| *Shewanella putrefaciens* 95 (CECT5346) | Isolated from putrefied butter | *Colección Española de Cultivos Tipo* (CECT) | Derby and Hammer, 1931 (31) | GCF_016406325.1 | YES |
| *Shewanella xiamenensis* HI32665 | Isolated from a coproculture | This study | This study | This study <sup>a</sup> | NO |

<sup>a</sup> Strain sequenced in this study.

The TYGS online service, which employs the same technique (i.e., GBDP) coupled with G+C content, provides a whole-genome-based GBPD tree phylogenetic analysis (25) (Figure 1). The tree was inferred with FastME from GBDP distances calculated from genome sequences obtained from the TYGS online service. The *S. seohaensis* strains, *S. xiamenensis* HI32665, and the type strain of *S. putrefaciens* and *S. decolorationis* were introduced to guarantee the representation of the genomes of interest. A duplicated entry means an automatic addition to the analysis by the TYGS service database. In the generated tree, DSM9451 clusters within *S. seohaensis* strains. These results further strengthen the suggestion that the correct classification of DSM9451 is *S. seohaensis*.

Although genome sequencing could identify DSM9451 as *S. seohaensis*, this method is not available in routine clinical microbiology laboratories as well as in many research laboratories. To explore alternative reliable methods of identification, a study focused on the phenotypic differentiation between *S. seohaensis, S. xiamenensis,* and *S. putrefaciens* was conducted.

### Phenotypic analysis provides incomplete discrimination between *Shewanella* species

To determine phenotypic differences between *Shewanella* species, the growth capacity of strains *S. seohaensis* DSM9451, *S. seohaensis* SA70, *S. seohaensis* CCUG60900, *S. putrefaciens* 95, and *S. xiamenensis* HI32665 at different temperatures, and their biochemical characterization were performed. The results were compared between the different strains and with those described in the literature for *S. decolorationis,* which has been used as a comparison species in previous studies (22, 26).

All strains of *S. seohaensis* exhibited growth at 37ᵒC, although they did not grow at temperatures of 4ᵒC or 42ᵒC (Supplementary Table 1). *S. putrefaciens* 95 showed poor growth at 37ᵒC but could grow at 4ᵒC. In contrast to what is described in the literature, all *S. seohaensis* strains and *S. xiamenensis* strains tested grew at 40ᵒC, indistinguishably from growth at the optimum temperature, as also described for *S. decolorationis* (22). Biochemical analysis revealed differences between strains of the same species as well as from phenotypes previously described in the literature for those strains. The KIA test revealed that CCUG60900 could produce hydrogen sulfide, contrary to what was described (26). In the API panels, divergences were observed within the *S. seohaensis* species (Supplementary Table 1). In the API ZYM panel and among the tested enzymes, alkaline phosphatases, butyrate esterase, caprylate esterase lipase, leucine aminopeptidase, α- chymotrypsin, acid phosphatase, and trypsin exhibited activity in all tested *S. seohaensis* strains. The trypsin activity observed on CCUG60900 contrasts with what was described previously (26). Notably, only DSM9451 presented enzymatic activity for N-acetyl-β-glucosaminidase. In the API 20NE panel, all strains tested positive for gelatine hydrolysis but displayed diverse responses regarding nitrate reduction: DSM9451 and CCUG 60900 tested positive, while SA70 tested negative (Supplementary Table 1). As for assimilation tests, all strains were able to use arabinose, N-acetyl-glucosamine, maltose, and malate as carbon sources. An exception occurred with the glucose utilization test, which was only positive for CCUG60900.

Overall, it is possible to phenotypically distinguish *S. seohaensis* from *S. putrefaciens* since the latter demonstrated the ability to grow at 4ᵒC, exhibit glucose fermentation in the KIA test, and does not hydrolyze gelatine or utilize arabinose and maltose as the carbon source in the API 20NE panel (Supplementary Table 1). In comparison to the phenotype described for *S. decolorationis*, these two species exhibit variations in leucine aminopeptidase activity and the assimilation of arabinose. Interestingly, the phenotype of *S. xiamenensis* closely resembled that of *S. seohaensis*. Our results revealed only minor discrepancies among these species, notably in the intensity of the signal for trypsin activity (negative) and esculin hydrolysis (positive). While other inconsistencies have been documented in the literature (Supplementary Table 1), we were unable to definitively confirm them in this study (22, 26).

### Whole-genome multilocus sequence typing scheme reveals species segregation and large strain diversity within species

For a more comprehensive understanding of the relationship between the isolates and to identify specific genes associated with each species, a wgMLST scheme was developed for all the strains analyzed in this study. For comparative analysis, closely related species including representatives of *S. decolorationis* and *S. xiamenensis*, were also incorporated. This scheme evaluates the diversity of the strains under study and identifies genes that are either present or absent in the various species. An exclusive gene was defined as being present in all isolates of a species and absent in isolates of all other species represented in the scheme. The scheme and the alleles identified are available in the chewie-NS (27) platform (https://chewbbaca.online/). Using the PHYLOViZ online software, it is possible to observe a minimum-spanning tree for the core-genome MLST (cgMLST) (Figure 2), i.e., of the allelic profiles of the genes present in all strains (cgMLST_100_ = 922 loci).

**Figure 2.**
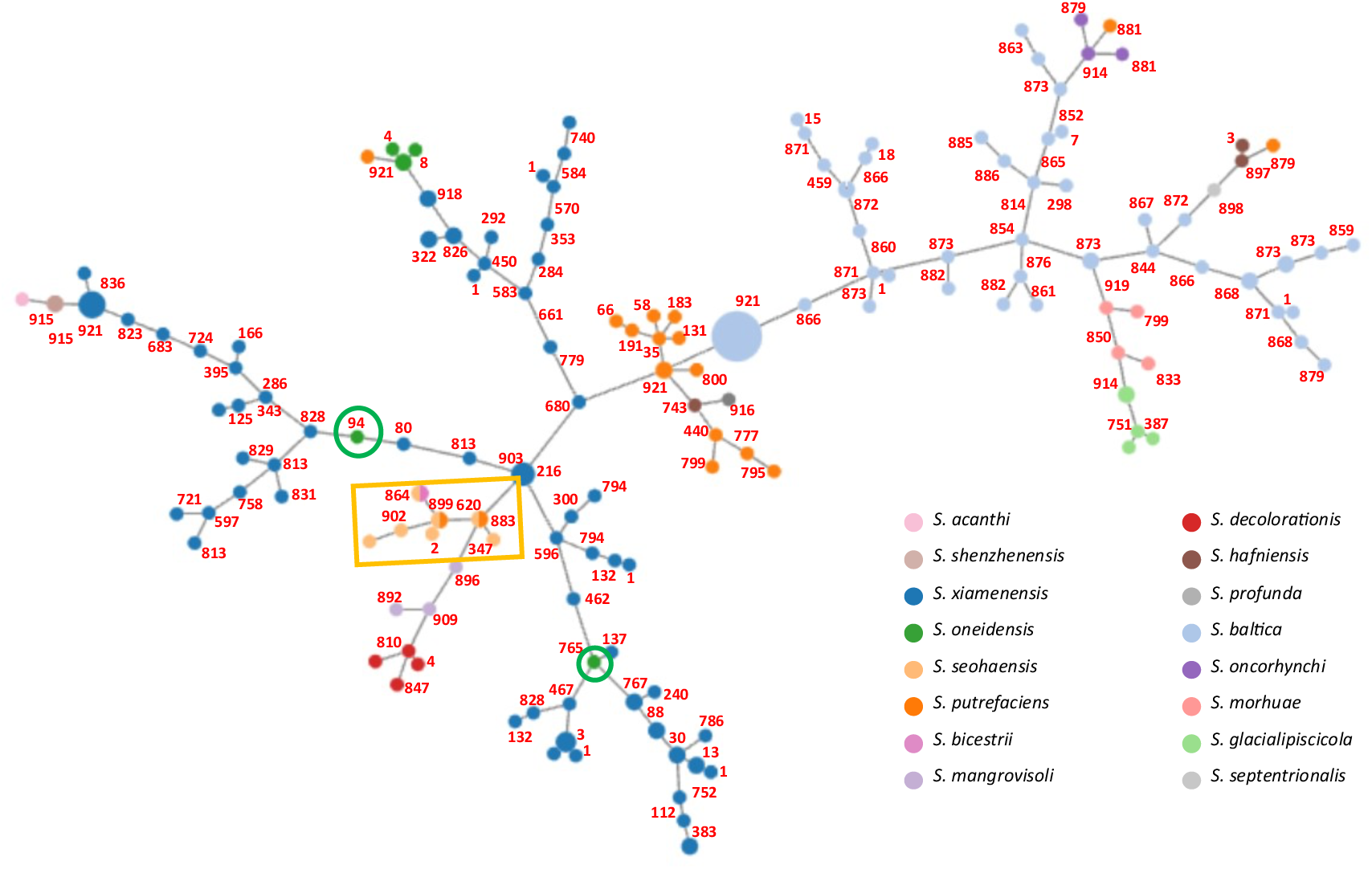
PHYLOViZ. The numbers represent the number of allelic differences between the two nodes. The green round box highlights the strains identified as *S. oneidensis* but in the *S. xiamenensis* cluster. The yellow rectangular box highlights the *S. seohaensis* strains.

In Figure 2 it is possible to observe sixteen different species, whose strains mostly segregate together in the tree. However, there are also several cases in which strains identified as a given species group with others of a different species. This is the case of the strains identified as *S. oneidensis* S2_009_000_R2_72 (GCA_003241225.1) and *S. oneidensis* SRR9109399 (GCA_945952185.1) that can be found among the *S. xiamenensis* cluster. Phylogenetic studies confirm they indeed belong to *S. xiamenensis* species (*d*DDH values are 76.50% [73.5 – 79.3%] and 71.20% [68.2 – 74%], respectively). The same situation could be observed with strains identified as *Shewanella putrefaciens*. In the *S. seohaensis* cluster (yellow rectangular box), *S. seohaensis* SA70 and *S. seohaensis* NCTC12093 have been misidentified as *S. putrefaciens* while *S. seohaensis* JAB-1 has been misidentified as *Shewanella bicestrii*, visible by the three circles with two colors in Figure 2.

A comparison of the genes of *S. seohaensis* with those of closely related species reveals a list of 8 exclusive genes (Supplementary Table 2). Interestingly, cgMLST analysis demonstrates extensive diversity within species, with the number of allelic differences between pairs of closest isolates of the same species being similar to pairs of closest isolates of different species (numbers on Figure 2). This phenomenon could explain the phenotypic differences observed within the same species. Additionally, without distinct phenotypic features for each species, phenotypic discrimination among them is not possible.

Given the absence of distinctive phenotypic characteristics between *S. seohaensis* and *S. xiamenensis,* this study allowed to identify exclusive genes that could function as targets for differentiating *S. seohaensis* from *S. xiamenensis* and other closely related species (see below).

### Novel method for the identification of *S. seohaensis* by detecting an exclusive gene

Among the 8 exclusive genes to *S. seohaensis* compared to closely related species in the previous section, the gene that encodes for a phosphatase PhoE (ID: VEE60752.1) was selected and corresponding primers were designed (Table 3 in Materials and Methods). A PCR was performed using genomic DNA from different strains, and the resulting agarose gel (Supplementary Figure 1) showed that amplification occurred only for the *S. seohaensis* strains. This confirmed the utility of this assay in identifying this species.

**Table 3.**
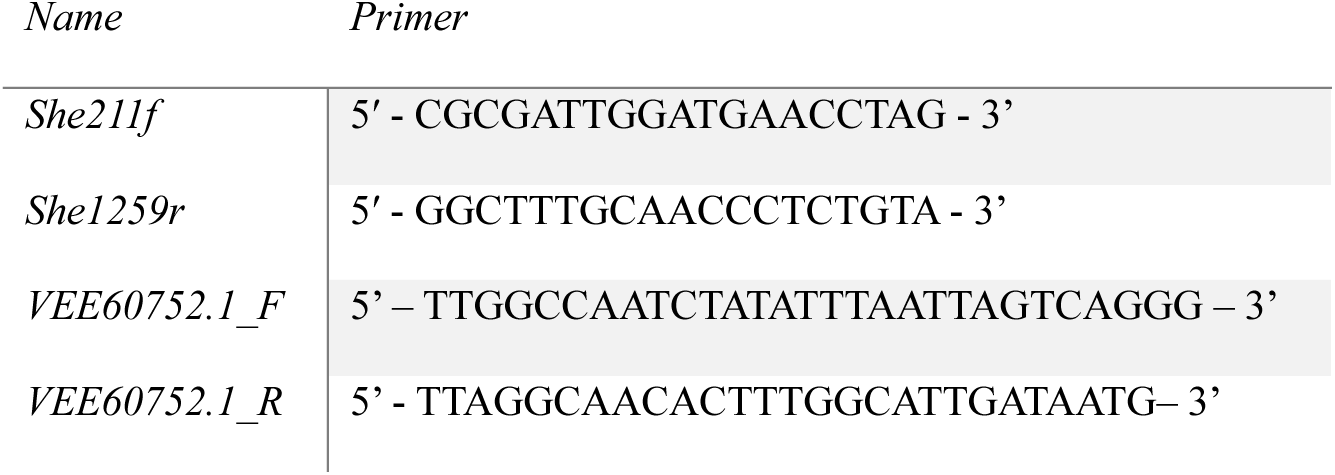
Primers used to confirm the presence of the exclusive genes.

These results provide a novel tool for distinguishing *S. seohaensis* from closely related species, including *S. xiamenensis* and *S. putrefaciens*, effectively resolving the issue of misidentification among these species.

## Conclusion

The misidentification of species within the *Shewanella* genus is well-documented (17, 18), especially between *S. seohaensis*, *S. putrefaciens,* and *S. xiamenensis* (21). *S. putrefaciens* often emerged as the initial taxonomic choice after preliminary tests (19, 24), which led to lumping together a highly diverse group of isolates. Moreover, the overall similarity between related species and the biochemical diversity within the same species increases the complexity of arriving at an accurate identification. Despite advances afforded by using MALDI-TOF for routine species identification in clinical microbiology laboratories, our results show that it still fails to accurately identify some species within the *Shewanella* genus, similar to what happens with other genera (28), possibly due to insufficient representation in existing databases. Although we identified several characteristics that can be used to distinguish *S. putrefaciens* from *S. seohaensis,* and *S. xiamenensis,* including the ability of these strains to grow at 4 ᵒC and ferment glucose in the KIA test, no phenotypic tests could reliably distinguish the latter two species. Distinguishing *S. seohaensis* from closely related species can be achieved by the PCR reaction proposed here, targeting a gene exclusive of *S. seohaensis*. Although we did not test strains representing all *Shewanella* spp. of the clade, *in silico* analysis indicates that this PCR reaction will also distinguish *S. seohaensis* from all other *Shewanella* genomes used to construct the wgMLST scheme proposed here.

Our work illustrates a way to overcome the limitations of several techniques for the identification of species within the *Shewanella* genus. While phenotypic traits may offer initial insights, complementing this information with the presence of species-specific genes, is an effective strategy for accurately identifying *Shewanella* isolates, serving as a practical and cost-effective alternative to genome sequencing. The wgMLST scheme proposed constitutes a framework allowing researchers to explore the diversity of each of these closely related species with potential consequences for our understanding of their pathogenic potential, environmental role, and gene flow between them. Several studies report *in vitro* experiments to obtain insights into the potential role of different *Shewanella* species in bioremediation processes (see for instance (29)). However, the large diversity observed within each of these species suggests that generalizing the results obtained from a single strain to the whole species may be particularly questionable in *Shewanella* spp. The wgMLST scheme proposed can help in providing a population context and help guide the choice of strains for further study. The growing recognition of the importance of *Shewanella* as potential infectious agents, in bioremediation, and in the environment at large highlights the relevance of accurate species identification to understand their interactions and specific roles in these processes.

## Materials and Methods

### Bacterial strains

The bacterial strains used in this work are described in Table 1. All strains were routinely cultivated on Luria-Bertani (LB) agar or in LB broth (30) at 37 ᵒC with aeration (at 150 rpm), except *S. putrefaciens* 95, which was cultured at 30 ᵒC.

The genomes of *S. putrefaciens* DSM9451 and *S. xiamenensis* HI32665 were sequenced to validate their identification, while the genomes of all other isolates are accessible on public databases.

### Genome sequencing

Genomic DNA from *S. putrefaciens* DSM9451 and *S. xiamenensis* HI32665 was extracted using the Invitrogen PureLink Genomic DNA mini kit (Thermo Fisher Scientific Inc., Waltham, MA, USA) according to the manufacturer’s instructions and using cultures grown in LB broth at 37 ᵒC overnight. The genomes were sequenced using an Illumina Nextseq® 500 system (2 X 150bp PE) (Illumina ©, San Diego, CA, USA), following the low-volume Nextera protocol (32). The reads were assembled with the INNuca pipeline v3.1 (33). For taxonomic classification, Kraken software 2.0.9 (34) was used.

### Phylogenetic analysis

The intergenomic distances were evaluated by digital DNA-DNA hybridization (*d*DDH) (35) and Average Nucleotide Identity (ANI) (36) methods. The technique used was Genome-BLAST Distance Phylogeny (GBDP), through the GGDC 3.0 online service (37), where the *d*DDH was calculated between the target genome and all the *Shewanella* genomes available in the database on 17^th^ June 2024. Strains of the same species are expected to exhibit dDDH values >70%, while isolates corresponding to the same subspecies should present values >79% (35). The values of *d*DDH were determined according to formula *d6*, which preserves most information (35). The TYGS online service, which employs the same technique (GBDP) combined with the G+C content, creates a whole-genome-based GBPD tree (25). ANI is a similar method to *d*DDH (36). An ANI value >95% indicates that the genomes belong to the same species (36, 38). Calculation of ANI values using the BLAST algorithm (ANIb) was achieved using JSpeciesWS v.4.1.1 (38).

### Phenotypic analysis

Temperatures permissive for growth were assessed by cultivating the different strains on LB agar at 4ᵒC, 30ᵒC, 37ᵒC, 40ᵒC, and 42ᵒC. The biochemical characterization of the different strains was performed following the protocols described in the “Clinical Microbiology Procedures Handbook” (39)and in line with the established standards for the *Shewanella* genus (4). Hydrogen sulfide production, fermentation of glucose and lactose, and gas production were tested in Kligler’s Iron Agar (KIA) (39). Enzymatic activities and carbon assimilation were tested using Analytical Profile Index (API) kits: API 20 NE and API ZYM (40–42). All phenotypic tests were performed in triplicate.

### Creation of a whole-genome multilocus sequence typing (wgMLST) scheme

To establish a finer relationship between isolates, a whole-genome multilocus sequence typing (wgMLST) scheme was developed. This was used to characterize the isolates sequenced in this study, as well as all genomes of closely related species of the *Shewanella* genus, including *S. baltica, S. putrefaciens, S. oneidensis, S. glacialipiscicola, S. morhuae, S. oncorhynchi, S. mangrovisoli, S. hafniensis, S. shenzhenensis, S. acanthi, S. profunda, S. septentrionalis, S. decolorationis* and *S. xiamenensis*, available on NCBI GenBank on the 17^th^ of June 2024. Supplementary Table 3 presents the list of all the assembly identifiers used in this study. Based on 202 draft or complete genomes, the wgMLST scheme was generated with chewBBACA v3.2.0 *CreateScheme* module (43), resulting in a scheme with 21046 loci representing the genetic diversity of the selected genomes. Upon verifying the quality of the allelic profiles, it was decided to remove 11 genomes due to poor quality (low representation of core genome loci) or unverified origin resulting in a final data set of 191 genomes. This data was visualized in PHYLOViZ online (44).

### Polymerase chain reaction for the identification of *S. seohaensis*

To differentiate *S. seohaensis* from closely related species, particularly the *S. putrefaciens* type strain and *S. xiamenensis*, one gene exclusive to *S. seohaensis* was selected, and specific primers were designed (VEE60752.1_F and _R – see Table 3). This gene (*phoE*), with a size of 609 bp, encodes for a phosphatase PhoE (ID: VEE60752.1). *In silico* PCR was performed to validate the amplification of a single product exclusively in the genomes of *S. seohaensis.* A PCR assay was performed for each strain using genomic DNA, the NZYTaq II 2x Green Master Mix (NZYTech, Lda, Lisboa, Portugal), and the primers indicated in Table 3. The 16sRNA gene, amplified with primers suitable for the *Shewanella* genus (She211f and She1259r) (45), was used as a control for identification as *Shewanella*.

## Data availability

To promote open science and reproducibility, all resources associated with this schema have been made publicly available. These include: the wgMLST schema and its annotations; the GenBank files used that can be used for functional annotation of loci; AlleleCall reports and AlleleCallEvaluator output; the wgMLST matrix of genetic profiles; and the metadata for all genomes used for schema creation and allele calling. The full dataset is available at: https://doi.org/10.5281/zenodo.13735156. The Schema and its annotations is available at: https://chewbbaca.online/species/.

## Acknowledgement

Financial support was provided by the following institutes and funding agencies with the respective projects specified: MOSTMICRO-ITQB base funding with references UIDB/04612/2020 and UIDP/04612/2020, LS4FUTURE Associated Laboratory (LA/P/0087/2020), Portuguese Foundation for Science and Technology (FCT) with the project grant PTDC/BIA-BQM/4143/2021 and the PhD grant 2022.12339.BD for Maria de Oliveira Firmino.

